# Antibacterial Activity of the Natural Product Elaiophylin against *Streptococcus suis* and its Biofilm Eradication Effect

**DOI:** 10.1101/2025.12.01.691497

**Authors:** Di Liu, Chang Chen, Qianqian Bao, Na Su, Yuhan Zhang, Xiliang Yang, Wei Fang, Chen Tan, Chenchen Wang, Manli Liu

## Abstract

The multidrug resistance (MDR) problem in *Streptococcus suis*(*S. suis*) is becoming increasingly severe, necessitating the development of novel antibacterial agents and strategies. In this study, seven Elaiophylin derivatives were isolated from Streptomyces sp. WS-30248, and the antibacterial activity and mechanism of action of the principal compound, Elaiophylin, against *S. suis* were systematically evaluated for the first time. Through comprehensive approaches including *in vitro* efficacy assays, biofilm inhibition and eradication tests, bacterial membrane integrity analysis, reactive oxygen species (ROS) level detection, and a mouse infection model, Elaiophylin was found to exhibit significant antibacterial activity against multiple clinically MDR *S. suis* strains. Its minimum inhibitory concentration (MIC) was as low as 0.5 μg/mL, and complete bactericidal activity was achieved within 24 hours at 4×MIC. The compound effectively inhibited and eradicated bacterial biofilms, directly killing embedded cells within the biofilm matrix. Mechanistic studies revealed that Elaiophylin functions through multiple synergistic pathways, including disruption of bacterial membrane integrity and induction of massive ROS accumulation, thereby interfering with the proton motive force (PMF), depleting intracellular ATP, and blocking energy metabolism. In the mouse infection model, the Elaiophylin-treated group showed a significantly increased survival rate of 60% and effectively reduced bacterial loads in tissues. This study demonstrates that Elaiophylin is a highly promising natural candidate drug with multi-target synergistic effects, offering a new strategy to combat MDR *S. suis* and biofilm-associated infections.

## 1. Introduction

Antimicrobial resistance (AMR) has been identified as a major global cause of mortality and has emerged as a core public health challenge of the 21st century. According to a 2019 World Health Organization (WHO) report, AMR was responsible for 700,000 deaths worldwide. It is estimated that by 2050, this death toll could rise to 20 million, with associated economic losses exceeding US $2.9 trillion. AMR has become a critical challenge, posing a serious threat to both human health and the global economic system [1–3]. Among the many pathogens exacerbating the antimicrobial resistance crisis, *S. suis*, a gram-positive bacterium with both high pathogenicity and zoonotic potential, not only serves as a key reservoir for AMR genes exhibiting high resistance to lincosamides, macrolides, and tetracyclines, but also possesses the ability to transfer these resistance genes to other pathogens. The evolution and dissemination of its resistance further underscore the complexity of this public health issue and intensify the AMR crisis [4–7] .*S. suis,* a type of zoonotic pathogen, can cause infections in both pigs and humans, leading to septicaemia and meningitis, and can even result in irreversible sequelae, including meningitis [8,9].China experienced two large-scale zoonotic outbreaks caused by *Streptococcus suis* serotype 2 (SS2) in 1998 and 2005. In the 1998 outbreak in Jiangsu, 25 people were infected, resulting in 14 deaths, and approximately 80,000 pigs were infected. In the 2005 outbreak in Sichuan, 215 human cases were reported, with 38 fatalities. Both outbreaks also led to extensive infection and mortality among pigs. The high pathogenicity of SS2 not only endangers life and health but also causes significant economic losses [10,11]. On the basis of the antigenic differences in capsular polysaccharides, 29 serotypes of *S. suis* have been identified to date. Strains of different serotypes exhibit marked heterogeneity in both their pathogenicity and resistance profiles. The efficacy of conventional antibiotics is compromised by resistance, while vaccines often struggle to achieve broad-spectrum coverage owing to serotype specificity [6,12–14]. In parallel, antivirulence strategies, such as targeting the pore-forming toxin suilysin, have emerged as promising alternative approaches to combat *S. suis* infections[15]. Therefore, there is an urgent need for drugs with broad-spectrum antibacterial activity and resistance potential against *S. suis*.

Natural products remain a primary source of new drugs and have long played a key role in the identification and development of antimicrobial agents, continuously providing novel chemical structures with highly potent antibacterial properties[16–18]. The inherent availability and rich chemical diversity of natural antibacterial products confer unique innate advantages in the antimicrobial field. Approximately two-thirds of clinically used antibacterial agents are derived from natural products. For decades, research on microbe-derived antibiotics through activity-guided purification has led to the identification of approximately 28,0 00 compounds, among which 200 have been found to be directly usable as drugs. Natural products continue to play a significant role in the development of new drugs and address the antibiotic crisis[19–23]. Theaflavins and their derivatives derived from black tea not only exhibit significant in vitro and in vivo antibacterial activity against SS2 but also exert antibacterial effects against *Bacillus coagulans* by adsorbing to surface phospholipids of cells, fully demonstrating the application potential of natural products in antimicrobial contexts[24,25]. Concurrently, previous work by our research group revealed that the natural product ellipticine not only has significant antibacterial activity against *S. suis*, effectively improving survival rates and reducing pathological damage in infected mice but also has been demonstrated to possess good in vitro and in vivo antibacterial activity against drug-resistant *Escherichia coli*, further highlighting the application value of natural products[26,27]. Therefore, in the context of the increasingly severe antimicrobial resistance of *S. suis*, there is an urgent need for novel chemical scaffolds, particularly the need to explore active molecules from natural product resources that can efficiently inhibit *S. suis*.

The sources of natural products are highly diverse and can be divided into three major categories: plant-derived, animal-derived, and microorganism-derived. Among microbial resources, actinomycetes are among the core synthetic sources of new antibiotics, with nearly two-thirds of known antibiotics produced by actinomycetes. Among antibiotics synthesized by actinomycetes, strains of the genus *Streptomyces* constitute the greatest proportion, accounting for approximately 80% of actinomycete-derived antibiotics[17,28–30]. Our research team has long been dedicated to the isolation of natural secondary metabolites. Through continuous exploration, we have isolated Elaiophylin and seven of its derivatives from the strain designated as WS-30248, which belongs to *Streptomyces philanthi*. Previous studies have confirmed that Elaiophylin and its derivatives exhibit broad-spectrum biological activities, including antibacterial, anthelmintic, anticancer, immunosuppressive, anti-inflammatory, and antiviral effects [31–35]. Currently, there have been no reports on the antibacterial activity or mechanism of Elaiophylin and its derivatives against *S. suis*. Therefore, in response to the clinical challenge of multi-serotype drug resistance in *S. suis*, this study aims to systematically investigate the antibacterial activity of these compounds against different serotypes of *S. suis*, their inhibitory and eradication effects on biofilm formation, and their underlying mechanisms. The findings are expected to provide a theoretical basis for the development of novel anti-*S. suis* agents, alleviate the crisis of antibiotic resistance, and address the current shortage of therapeutic options for infections caused by drug-resistant *S. suis*.

## 2. Results

### 2.1 Antibacterial Activity of Elaiophylin Derivatives

To identify the antibacterial constituents of the isolated strain WS-30248, we first observed its colonial morphology. The MIC values of compounds 1–7 and tetracycline against various bacterial strains were determined (**Table 1**). The results revealed that these derivatives exhibited significant inhibitory effects against Gram-positive bacteria, whereas their MIC values against Gram-negative bacteria all exceeded 128 μg/mL, indicating a distinct selective antibacterial activity toward Gram-positive pathogens. Notably, compound 4 (with a methyl group at R₁ and hydrogen at R₂–R₅) demonstrated particularly potent activity against *S. suis* SC19, with an MIC value as low as 0.5 μg/mL, representing the strongest anti-*S. suis* activity among all tested compounds. Furthermore, compounds 1, 2, 4, 6, and 7 all showed MIC values of 0.5 μg/mL against *S. aureus* ATCC 25923. In contrast, compound 3, in which all R₁–R₅ positions are hydrogen-substituted, exhibited markedly reduced antibacterial activity, with an MIC of 1 μg/mL against the same strain. These findings suggest that the antibacterial potency of Elaiophylin derivatives is closely associated with the type and position of substitutions at the R₁–R₅ sites. The presence of methyl versus hydrogen substituents critically influences the activity level. Given its outstanding inhibitory efficacy against *S. suis* SC19, compound 4 (Elaiophylin) was selected as the primary focus for subsequent investigation. Further studies will concentrate on elucidating its antibacterial activity and mechanism of action against this pathogen, thereby providing a theoretical foundation for future drug development.

**Table 1.**
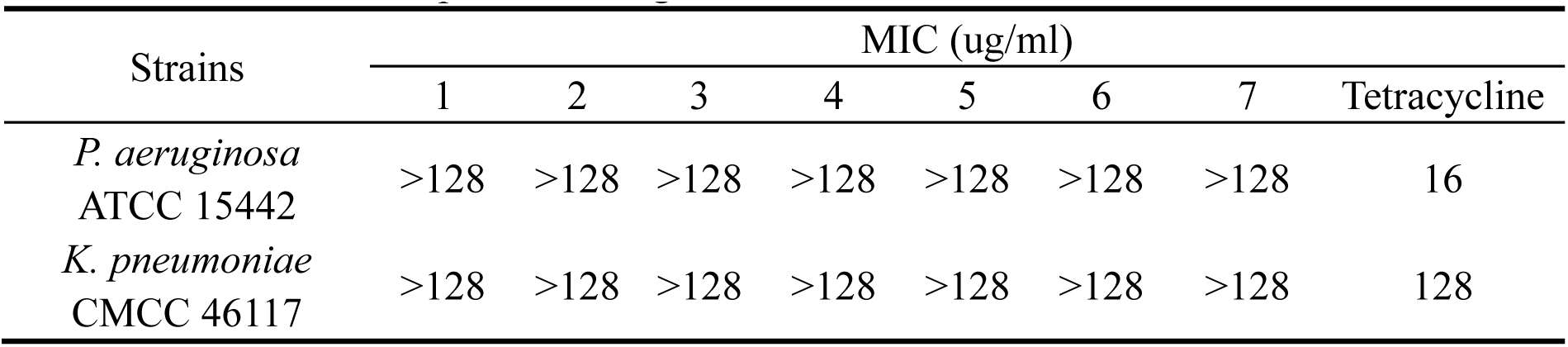
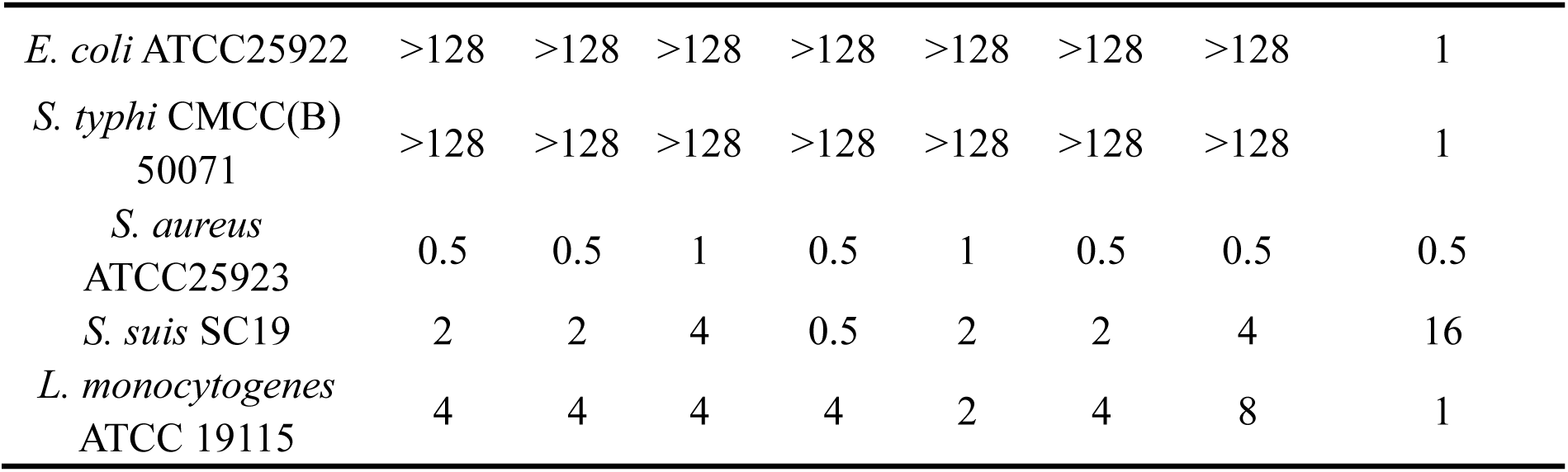
MICs of compounds 1-7 against various bacterial strains.

### 2.2 Antibacterial Activity of Elaiophylin against Clinically Resistant *S. suis*

The bacteriostatic and bactericidal activities of Elaiophylin against five different serotypes of *S. suis* were determined via MIC and MBC assays using clinically isolated strains, with five strains of each serotype selected for testing. As shown in **Table 2**, the MIC values of Elaiophylin against these strains ranged from 0.5 to 2 μg/mL, and the MBC values ranged from 1 to 4 μg/mL. Elaiophylin exhibited significant and stable antibacterial and bactericidal activities against different serotypes of *S. suis*, highlighting its advantages in covering multiple bacterial strains. These results indicate that Elaiophylin is an effective candidate antibiotic against multiple serotypes of *S. suis*.

**Table 2.**
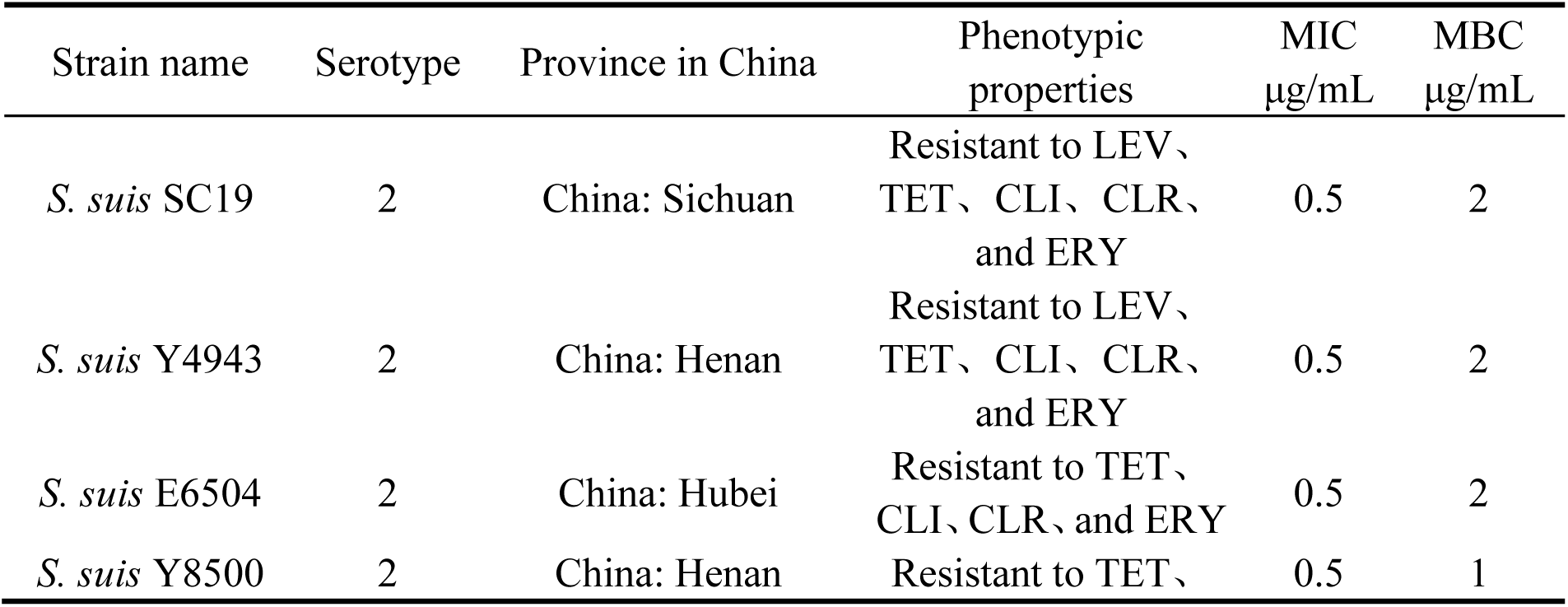
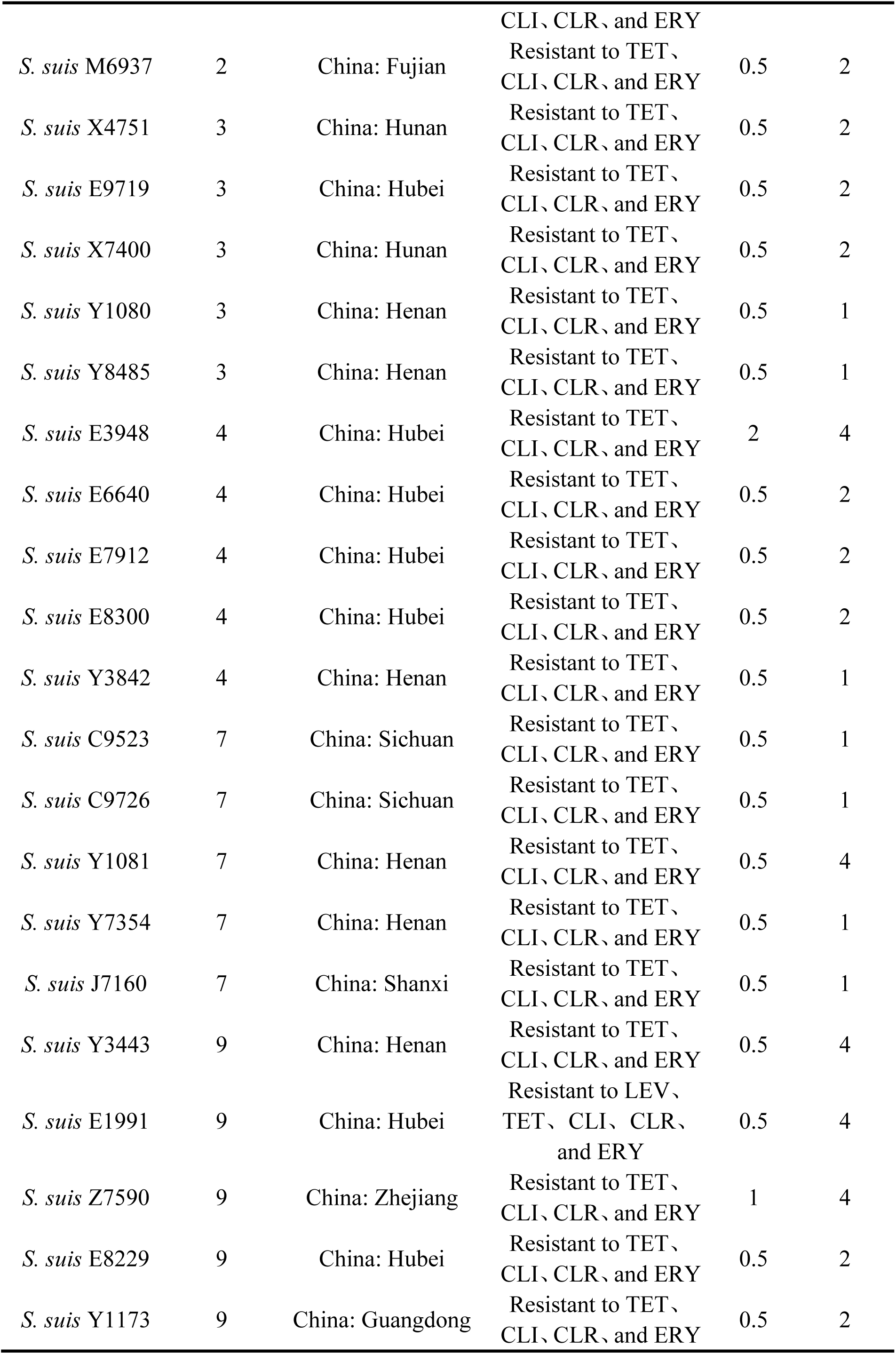
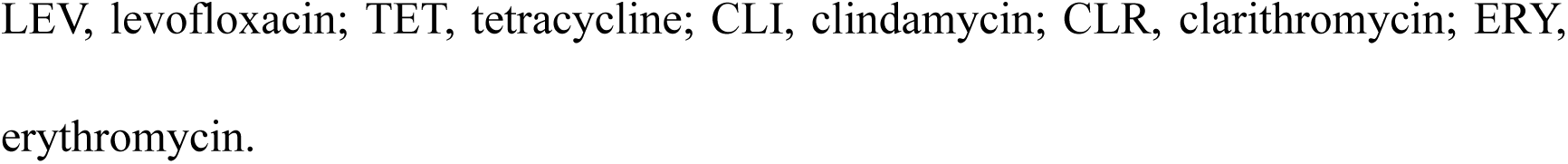
MICs and MBCs of Elaiophylin against different serotypes of *S. suis*.

### 2.3 Elaiophylin effectively kills virulent strains of *S. suis*

Based on the potent anti-*S. suis* activity of Elaiophylin against strain SC19, we further sought to systematically evaluate its antibacterial efficacy across various serotypes of *S. suis*. To further evaluate the antibacterial activity of Elaiophylin, five strains of different serotypes were selected for the systematic determination of growth curves and time‒kill curves to investigate its antibacterial effects across multiple strains. The results demonstrated that both 2×MIC and 4×MIC achieved complete inhibition of all five different serotypes of *S. suis*. In contrast to the continuous proliferation trend observed in the control group, these two concentrations completely suppressed the growth of the five different serotype strains (**Fig. 2A-E**). This finding indicates that Elaiophylin exerts potent antibacterial activity against multiple serotypes even at lower concentrations (2×MIC), whereas the inhibitory effect remains equally stable at higher concentrations (4×MIC), highlighting its broad-spectrum coverage capability against different serotypes of *S. suis*. The bactericidal efficacy was further investigated through time‒kill curves. At 2×MIC, the viable bacterial counts of all five different serotypes of *S. suis* were significantly lower than those in the control group. The 4×MIC group achieved complete bactericidal activity against these five strains within 24 hours, with no signs of bacterial regrowth observed throughout the entire culture period (**Fig. 2. Antibacterial activity of**). These findings demonstrate that Elaiophylin possesses potent and sustained bactericidal activity, particularly exhibiting stable efficacy against multiple serotypes, providing key evidence for subsequent validation of its potential anti-resistance applications.

**Fig. 1.**
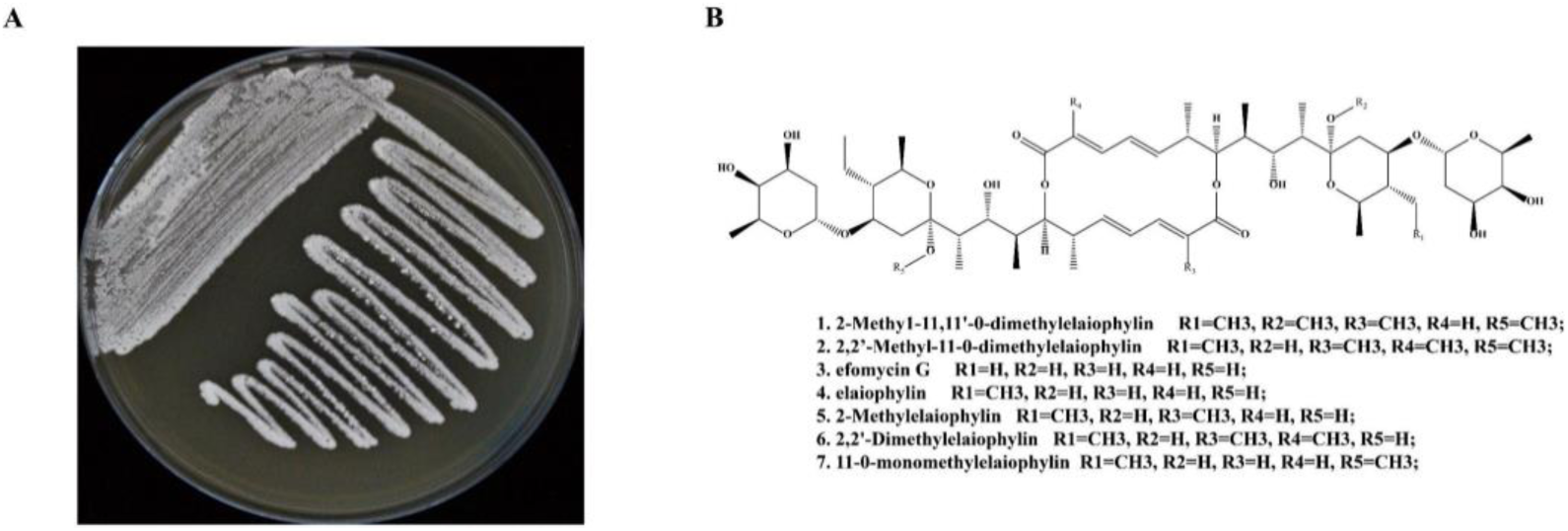
Morphology of the isolate WS-30248 and structures of the isolated compounds. (A) Colonial morphology of strain WS-30248. (B) Structures of compounds 1-7.

**Fig. 2.**
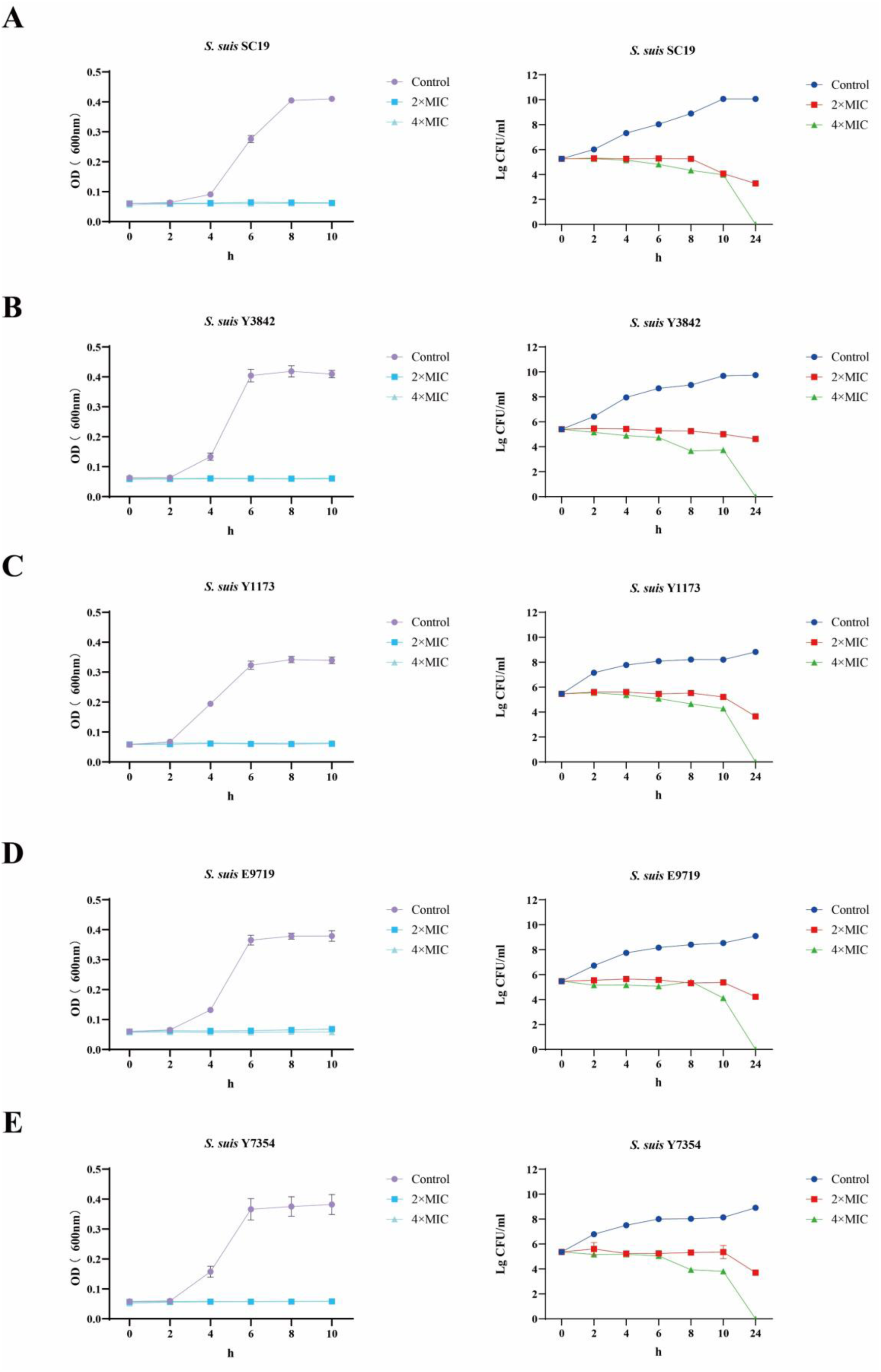
Antibacterial activity of Elaiophylin against five different serotypes of *S. suis*. A: Growth curves and time-kill curves of S. suis strain SC19; B: Growth curves and time-kill curves of S. suis strain Y3842; C: Growth curves and time-kill curves of S. suis strain Y1173; D: Growth curves and time-kill curves of S. suis strain E9719; E: Growth curves and time-kill curves of S. suis strain Y7354.

### 2.4 Elaiophylin Effectively Combats Bacterial Biofilm and Inhibits Biofilm Formation

Bacterial biofilms are key inducers of chronic and persistent infections, posing a severe challenge to clinical anti-infective therapy.[36,37] This study further investigated the inhibitory and eradicating effects of Elaiophylin on *S. suis* biofilms. *S. suis* SC19, a highly pathogenic, multidrug-resistant, and biofilm-forming dominant epidemic strain of *S. suis*, was selected for biofilm experiments. The anti-biofilm effect of Elaiophylin against S. suis SC19 was first visualized by crystal violet staining, which revealed a clear reduction in biofilm mass. Quantitative assessment further established a significant, concentration-dependent inhibition of biofilm formation by Elaiophylin(**Fig. 3A**), reaching an inhibition rate of over 89.5% at 4 μg/mL Elaiophylin. No viable bacteria were detected in the biofilms of the 4 μg/mL group, confirming that Elaiophylin effectively inhibited bacterial proliferation within the biofilms (**Fig. 3B, C**). Additionally, biofilm eradication experiments (Figure D) revealed that Elaiophylin also has concentration-dependent eradication activity against mature biofilms, with an eradication rate of over 75.8% at 4 μg/mL Elaiophylin, significantly reducing the biomass of mature biofilms. Moreover, 4 μg/mL Elaiophylin reduced the viable bacterial count within the biofilm by 3 log values (**Fig. 3E, F**). To visually evaluate the impact of Elaiophylin on bacterial activity within *S. suis* biofilms, this study used Syto Green (a live-cell dye) and propidium iodide (PI, a dead-cell dye) for live/dead staining of the biofilms. Observations via confocal laser scanning microscopy revealed that in the untreated control group, the green fluorescence was dense and uniform, whereas the red fluorescence was extremely weak, indicating that the biofilm was predominantly composed of live bacteria. As the concentration gradient of Elaiophylin increased, red fluorescence signals gradually intensified in the biofilm, whereas green fluorescence correspondingly weakened. In particular, at concentrations of 2 μg/mL and above, red fluorescence was widely distributed, and green fluorescence appeared fragmented and sparse. This result visually confirms that Elaiophylin effectively penetrates the *S. suis* biofilm matrix, significantly increases bacterial mortality within the biofilm, and potently kills live bacteria **(Fig. 4)**. In summary, Elaiophylin clearly inhibits and eradicates S. suis SC19 biofilms in a concentration-dependent manner, and its dual-action characteristics provide a potential strategy for overcoming the challenges of chronic infection and drug resistance mediated by *S. suis* biofilms.

**Fig. 3.**
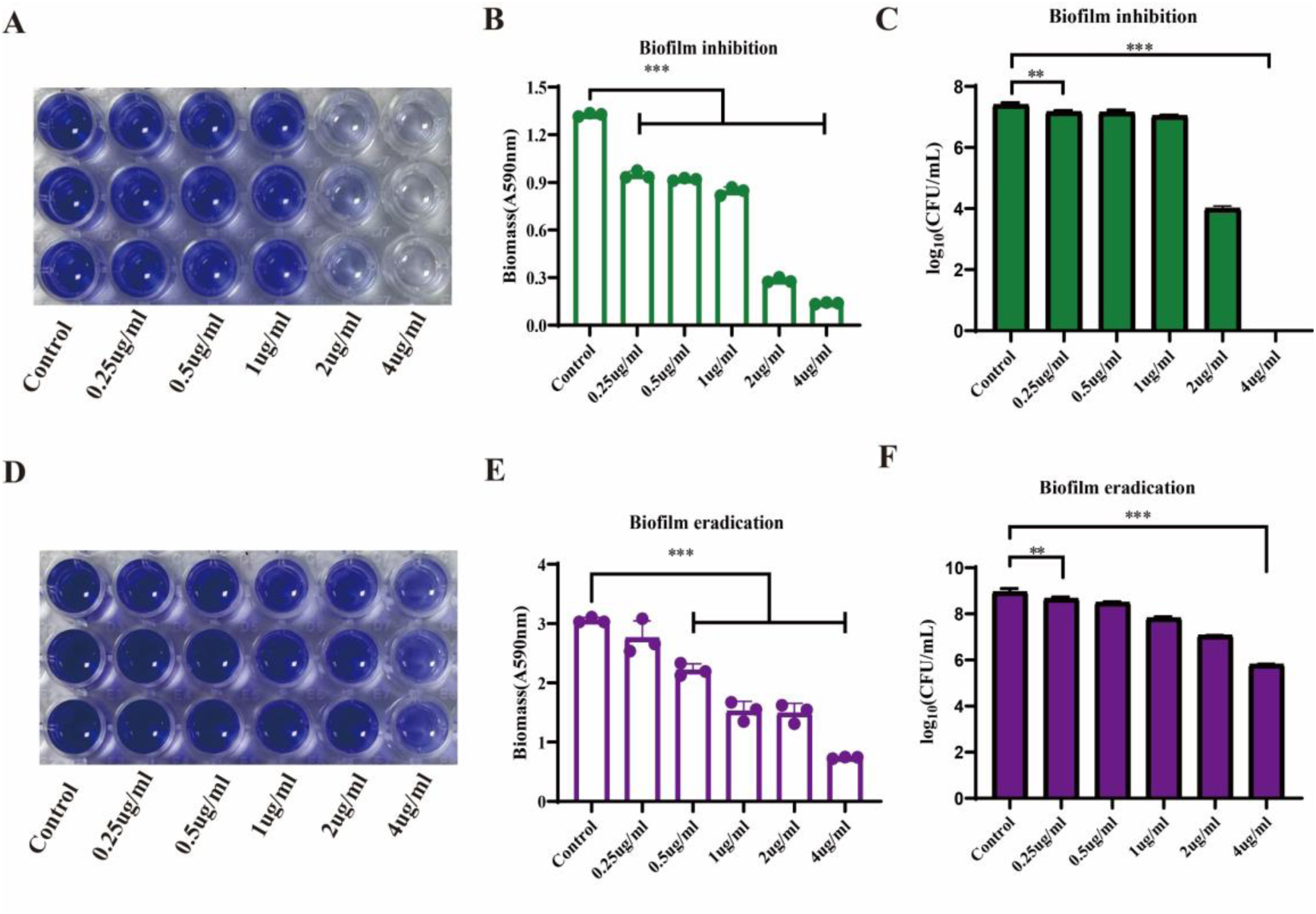
Effects of different concentrations of Elaiophylin on *S. suis* SC19 biofilms. (A, B) The inhibitory effect of Elaiophylin on bacterial biofilms was evaluated via CV staining. (C) Bactericidal effect of Elaiophylin on bacterial biofilm inhibition. (D, E) The eradicating effect of Elaiophylin on bacterial biofilms was evaluated via CV staining. (F) Bactericidal effect of Elaiophylin on bacterial biofilm eradication. (The data are presented as the means ± SDs of three biological replicates, ** <0.01, *** <0.001).

**Fig. 4.**
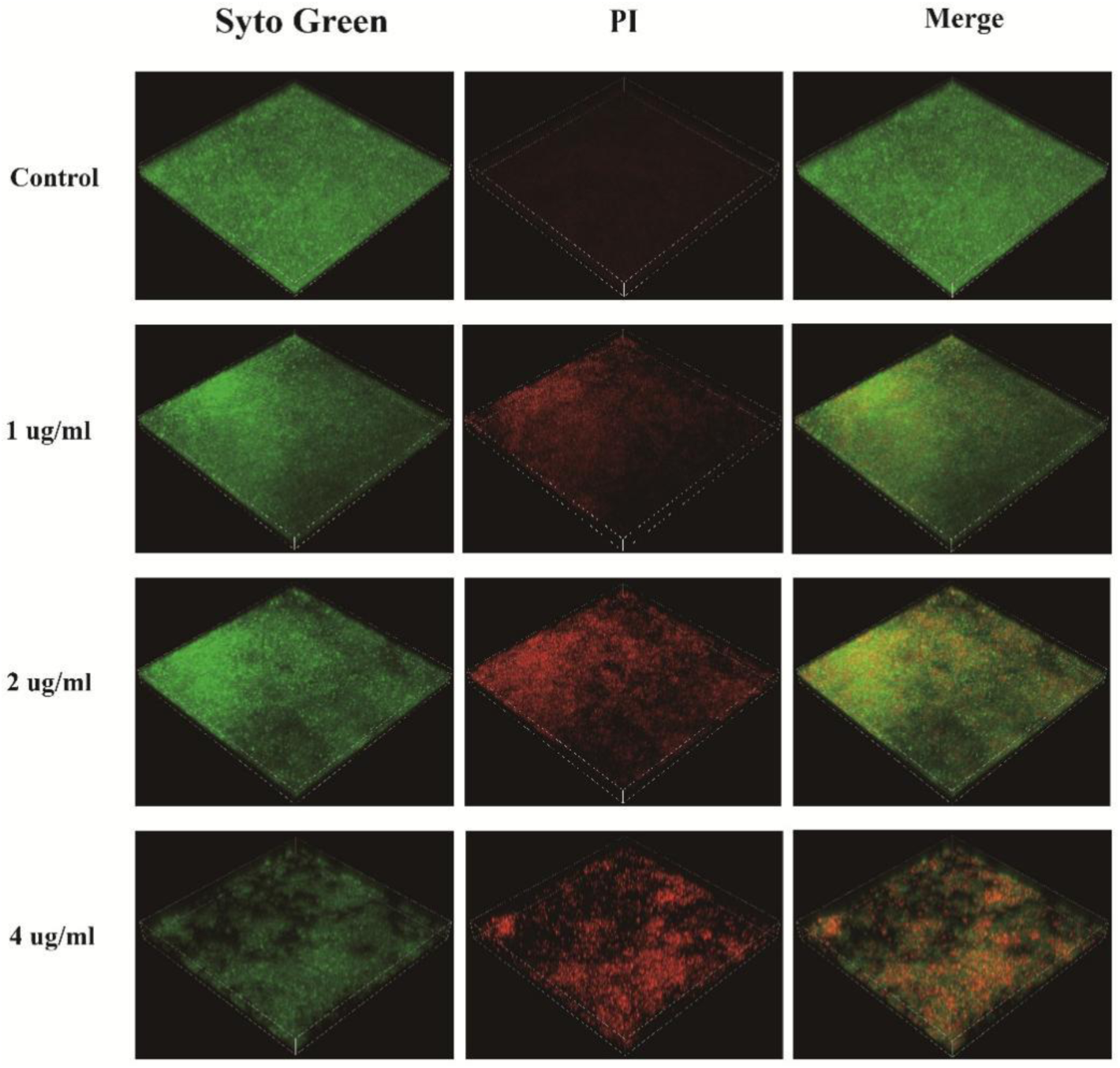
Live/dead bacterial distribution in *S. suis* SC19 biofilms treated with different concentrations of Elaiophylin, as observed via confocal laser scanning microscopy with SYTO Green and PI staining.

### 2.5 Membrane-Damaging Effect of Elaiophylin and the Induction of Bacterial Oxidative Stress

To investigate the antibacterial mechanism of Elaiophylin against *S. suis*, this study assessed its effect on the cytoplasmic membrane of strain SC19 via the dye propidium iodide (PI). Compared with the control treatment, Elaiophylin treatment resulted in a significant increase in the fluorescence intensity of *S. suis* in a dose-dependent manner, indicating its ability to disrupt bacterial membrane integrity (**Fig. 5A**). The results of ROS level detection revealed that the fluorescence intensity in the Elaiophylin-treated group also increased significantly, demonstrating that the drug can induce a strong oxidative stress response in bacteria, accelerating membrane damage and bacterial death (**Fig. 5B**). To further explore its interference mechanism with the core driving force of bacterial energy metabolism, the effects on the proton motive force (PMF) were investigated via DiSC₃(5) and BCECF-AM probes. The fluorescence of DiSC₃(5) decreased significantly with increasing drug concentration, suggesting membrane potential depolarization (**Fig. 5C, D**); the increase in BCECF-AM fluorescence indicated disruption of the proton gradient. The PMF, which is composed of the membrane potential and proton gradient, serves as the core driver of energy metabolism and the driving force for ATP synthesis. Therefore, we further examined the effect of Elaiophylin on bacterial ATP levels. Intracellular ATP measurements revealed a significant dose-dependent decrease in ATP content as the concentration of Elaiophylin increased (**Fig. 5E, F**). Thus, the drug exerts its antibacterial effect by disrupting PMF and consequently interfering with energy metabolism. The extracellular ATP results revealed a significant increase in ATP leakage, further suggesting that Elaiophylin enhances cell membrane permeability, leading to energy loss. Given that the PMF is essential for driving flagellar-mediated motility in bacteria, we further assessed the effect of Elaiophylin on the motility of *S. suis* SC19 using a swimming assay. The results demonstrated a marked inhibition of bacterial migration in the treated group, with a significantly reduced swimming zone diameter compared to the control. In conclusion, Elaiophylin exerts its antibacterial effect against *S. suis* by disrupting cell membrane integrity and interfering with the PMF.

**Fig. 5.**
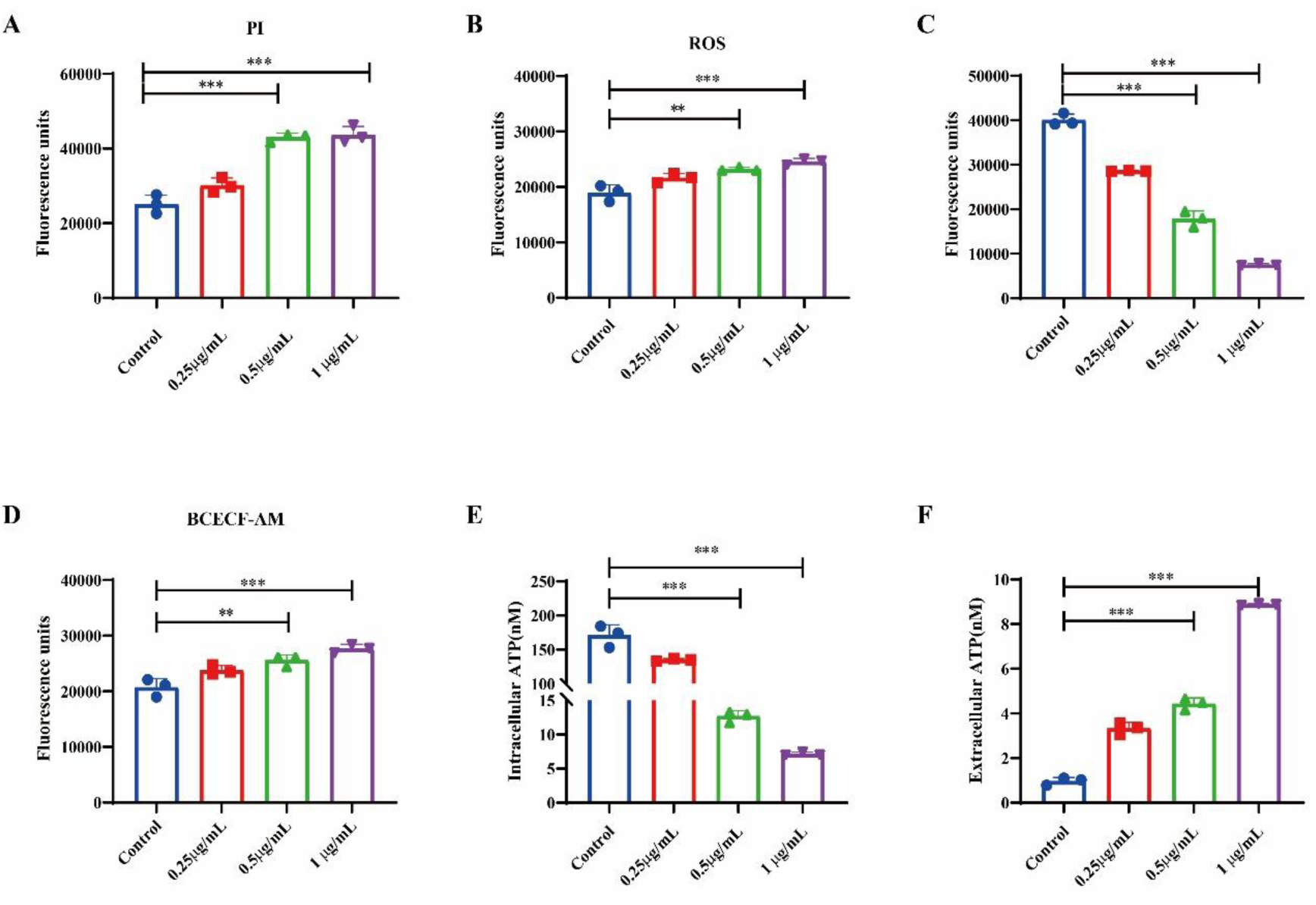
Effects of different concentrations of Elaiophylin on cell membrane integrity, oxidative stress, proton motive force, and energy metabolism in the SC19 strain. A: Cell membrane integrity detected by the PI fluorescent probe; B: Reactive oxygen species level detection; C: Membrane potential detected by the DiSC₃(5) fluorescent probe; D: Transmembrane proton gradient detected by the BCECF-AM probe; E: Intracellular ATP content measurement; F: Extracellular ATP leakage detection. (The data are presented as the means ± SDs of three biological replicates, **<0.01, ***<0.001).

**Fig. 6.**
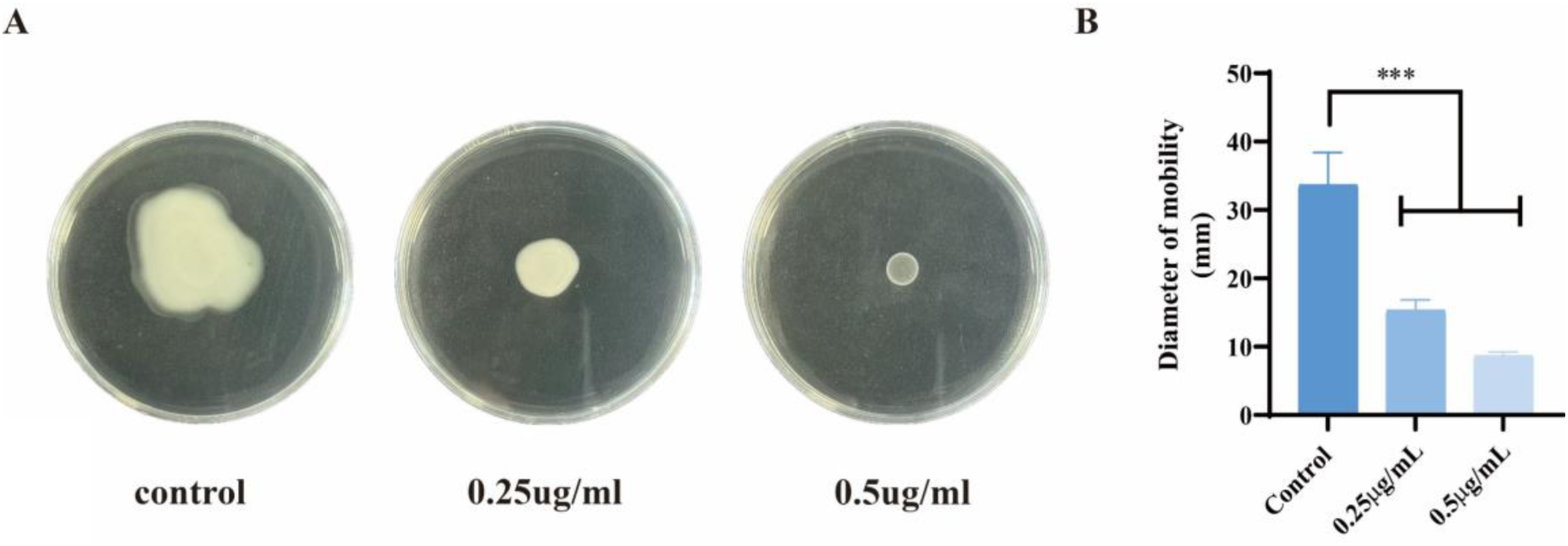
Effect of Elaiophylin on the motility of *S. suis* SC19 on agar plates. (A) Representative image of the motility zone of *S. suis* SC19. (B) Diameter of the bacterial motility zone. (The data are presented as the means ± SDs of three biological replicates; ***<0.01; ns indicates no statistical significance).

**Fig. 7.**
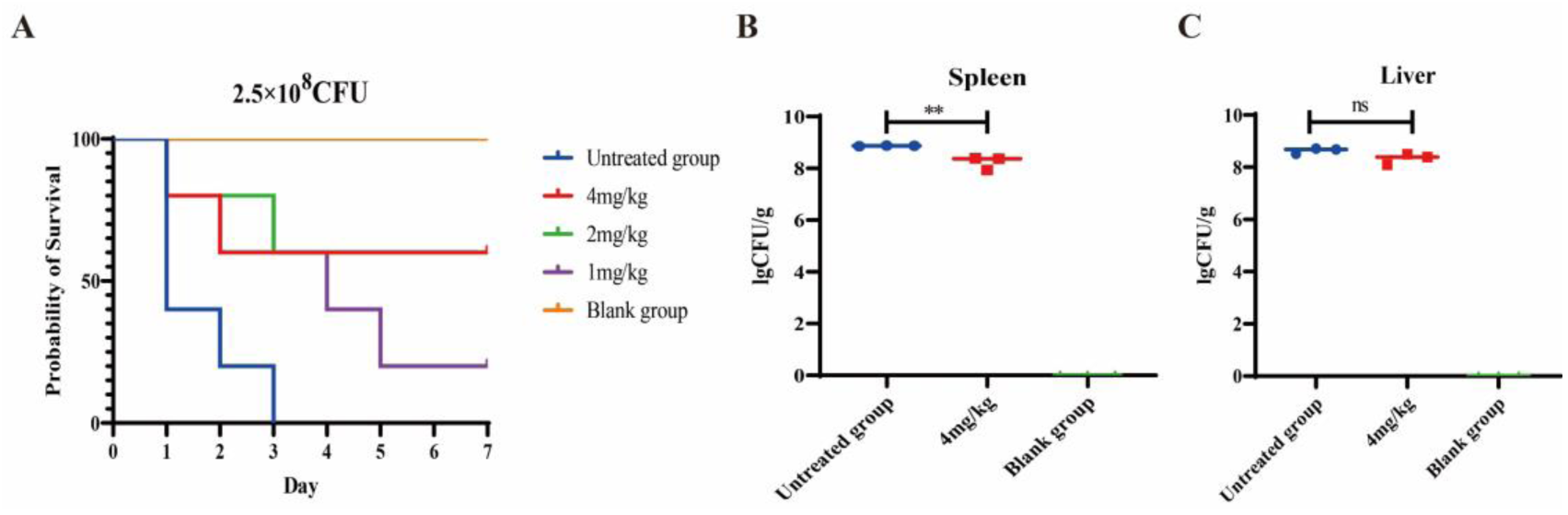
Elaiophylin demonstrates therapeutic efficacy in a mouse model of peritonitis infection. (A) Infected mice were treated with the drug 1 hour post-infection. (B, C) Effect of the drug on the bacterial load in the spleen and liver of infected mice. (The data are presented as the means ± SDs of three biological replicates; **<0.01; ns indicates no statistical significance).

### 2.6 In Vivo Antibacterial Activity of Elaiophylin

To determine the in vivo antibacterial efficacy of Elaiophylin, we established a mouse model of acute peritonitis infection and observed the survival rates of the mice in different treatment groups. The results revealed that the untreated group reached 100% mortality within 3 days post-infection. All the Elaiophylin treatment groups (4 mg/kg, 2 mg/kg, and 1 mg/kg) exhibited significant therapeutic protective effects, with survival rates reaching 60% in the 4 mg/kg and 2 mg/kg dose groups. (). Furthermore, we performed quantitative analysis of bacterial colonization in the spleen and liver tissues of infected mice to evaluate the in vivo bacterial clearance effect of Elaiophylin. The results revealed high bacterial loads (lgCFU/g) of *S. suis* SC19 in the spleen and liver tissues of the untreated group. After treatment with 4 mg/kg Elaiophylin, the bacterial load in the spleen tissue was significantly reduced, indicating that Elaiophylin effectively clears *S. suis* in the spleen. Although no statistically significant difference was observed in the liver tissue, the bacterial load showed an overall decreasing trend (). These results visually demonstrate the clearance effect of Elaiophylin against *S. suis* in vivo and fully confirm its good in vivo therapeutic efficacy, laying an experimental foundation for its subsequent clinical translation and application in animal models.

## 3. Discussion

*S. suis* is not only one of the most common pathogenic microorganisms in large-scale pig farms but also a significant pathogen capable of inducing severe infectious diseases in humans, posing a substantial threat to public health security [38,39]. Currently, the development of new antibiotics lags far behind the evolution of bacterial resistance, urgently necessitating the exploration of novel drug scaffolds to address this critical challenge[40,41]. Natural products have long served as a cornerstone for the development of antimicrobial agents, with those derived from Streptomyces playing a particularly prominent role. In this study, seven Elaiophylin derivatives purified from Streptomyces sp. WS-30248 demonstrated distinct and selective antibacterial activity against Gram-positive bacteria, including *S. aureus*, *S. suis*, and *L. monocytogenes*. Notably, Elaiophylin exhibited the most potent inhibitory activity against *S. suis* SC19. As a representative bioactive natural product, Elaiophylin belongs to the family of C2-symmetric glycosylated 16-membered macrolides, which have been confirmed to possess unique and broad pharmacological properties [35,42]. The characteristic structural flexibility and bioactivity adaptability of its macrolide scaffold offer a potential strategy to address the challenge of multi-serotype heterogeneity in *S. suis* and the limited efficacy of conventional antibiotics. Through systematic investigation, this study revealed that Elaiophylin not only exhibits robust antibacterial activity across diverse *S. suis* serotypes but also demonstrates biofilm-interfering effects, thereby providing key experimental evidence to advance research on natural product-based strategies against drug-resistant bacteria and to help mitigate the ongoing antibiotic resistance crisis.

Elaiophylin has significant and stable bacteriostatic and bactericidal effects on multiple serotypes of clinically isolated *S. suis*. Notably, the multidrug-resistant strains SC19, Y4943, and E1991 are resistant to five commonly used antibiotics, including the fluoroquinolone levofloxacin, the tetracycline tetracycline, the macrolides erythromycin and clarithromycin, and the lincosamide clindamycin. Among these, the MIC of erythromycin against strain Y4943 is greater than 128 μg/mL [43]. Although both Elaiophylin and erythromycin are macrolides, Elaiophylin exhibits excellent antibacterial activity against Y4943, with an MIC of only 0.5 μg/mL and an MBC of 2–4 μg/mL, fully demonstrating the potential application value of Elaiophylin against multidrug-resistant *S. suis*. Elaiophylin at 2×MIC completely inhibits the growth of multiserotype *S. suis*, and at 4×MIC, it achieves complete bactericidal activity within 24 hours without bacterial regrowth. The rapid and sustained antibacterial kinetic characteristics of *S. suis* provide potential for addressing acute *S. suis* infections and developing broad-spectrum anti-*S. suis* drugs. Bacterial biofilms not only significantly increase tolerance to various antibiotics and disinfectants but also readily cause refractory chronic infections, posing enormous challenges to clinical anti-infective therapy [44,45]. In this study, Elaiophylin effectively inhibited and eradicated *S. suis* biofilms and demonstrated the ability to efficiently penetrate the biofilm matrix and kill embedded bacteria, offering considerable application prospects for overcoming biofilm-mediated resistance and intervening in persistent biofilm infections.

At the mechanistic level, Elaiophylin treatment significantly induced massive ROS generation in *S. suis*, indicating that oxidative stress plays a key role in the bactericidal activity of Elaiophylin against *S. suis*. This finding is consistent with the conclusions of Kohanski et al. (2007), who emphasized the critical role of oxidative stress in antibacterial agent-induced bacterial death[46]. Grigoriev et al. (2001) reported that Elaiophylin can form stable conductive ion channels in the bacterial cell membrane, significantly disrupting cellular ion gradients and potential states and ultimately inhibiting bacterial growth and reproduction [47]. The membrane damage effect of Elaiophylin on bacteria aligns precisely with this mechanism. Concurrently, the synergistic mode of action between this membrane damage effect and the disruption of energy metabolism enables it to efficiently penetrate the *S. suis* biofilm matrix and kill bacteria within the biofilm. This finding correlates with the observed phenotype of biofilm inhibition and eradication, validating the consistency between the mechanism of action and functional manifestation. Furthermore, combined with the survival data from the mouse acute peritonitis infection animal model, all the Elaiophylin treatment groups showed significant therapeutic effects on infected mice. The 4 mg/kg and 2 mg/kg dose groups achieved a survival rate of 60%, which intuitively verifies the in vivo antibacterial efficacy from an in vivo perspective, providing robust in vivo experimental evidence for its subsequent development as a candidate drug against *S. suis* infection. Future studies should further explore its pharmacokinetics and safety in animal models and further elucidate its molecular targets to promote its clinical translation and expand its clinical application in anti-resistant bacterial therapy.

In summary, Elaiophylin has significant efficacy against multiserotype, multidrug-resistant *S. suis* and biofilm infections. Through a multipathway synergistic mode involving membrane disruption, the induction of oxidative stress, and interference with energy metabolism, it achieves rapid bactericidal action, complete growth inhibition, and efficient biofilm eradication in vitro, demonstrating substantial antibacterial potential. This provides a new candidate drug and intervention strategy for treating drug-resistant *S. suis* infections. This study not only offers new insights for addressing the challenges of *S. suis* resistance and biofilms but also may serve as a reference for treating other drug-resistant gram-positive bacterial infections, thereby further expanding the application scope of natural products in antibacterial therapy.

## 4. Materials and methods

### 4.1 Bacteria and Reagents

The clinical strains used in this study were obtained from laboratory isolates and preserved strains, including *S. suis* SC19 and other clinical strains. These strains were routinely cultured in Tryptic Soy Broth (TSB; BD, MD Rockville, USA) liquid media or on Tryptic Soy Agar (TSA; BD, MD Rockville, USA) solid plates supplemented with an appropriate concentration of newborn calf serum (China Four Seasons Green Company) at 37°C. For antimicrobial susceptibility testing, Mueller–Hinton broth (MHB, Invitrogen, USA) was used. The compounds were isolated by our laboratory.

### 4.2 Antibacterial Assays

The minimum inhibitory concentration (MIC) was determined via the microbroth dilution method in accordance with the guidelines of the Clinical and Laboratory Standards Institute (CLSI). The procedure was as follows: the bacterial stock was streaked onto TSA solid medium containing 5% newborn calf serum and incubated overnight at 37°C. The cultured bacterial mixture was then subcultured at a 1:100 ratio into fresh TSB liquid medium supplemented with 10% newborn calf serum. After cultivation, the bacteria were collected via centrifugation, resuspended, and adjusted to an OD600 of 1.0, followed by dilution to a final concentration of 5×10⁵ CFU/mL. For each drug and bacterial strain, three replicate wells were set up, along with negative controls (medium only) and positive controls. The MIC values were determined by observing bacterial growth. After the 16-h incubation endpoint for MIC determination, the 96-well cell culture plate was removed. Each well was vortexed and mixed thoroughly, and the bacterial suspension was spread onto TSA solid medium containing 5% newborn calf serum. After incubation at 37°C, the minimum bactericidal concentration (MBC) was determined. Three replicates were set for each group.

### 4.3 Growth Curve Testing

Bacterial culture was performed in a 96-well plate. Each well received 100 μL of the drug at final concentrations of 2×MIC or 4×MIC and 100 μL of the bacterial suspension. The plate was incubated at 37°C for 10 hours, and OD₆₀₀ measurements were taken every 2 hours to assess streptococcal growth.

### 4.4 Time‒Kill Curve

Referring to Section 4.4. During the incubation period, 100 μL of the mixture was collected every 2 hours, subjected to 10-fold serial dilution, and then plated onto TSA solid medium containing 5% newborn calf serum. After incubation, the data were recorded, and a time‒kill curve was plotted.

### 4.5 Biofilm assay

Biofilms were prepared in 96-well plates. Each well received 100 μL of diluted bacterial suspension and 100 μL of TSB medium containing the drug, which were mixed thoroughly. Negative control wells containing only 200 μL of TSB medium and positive control wells containing only 200 μL of bacterial suspension were also set up. After 24 hours of cultivation, the planktonic cells were aspirated and discarded. The wells were gently washed three times with PBS buffer, the PBS was discarded, and 10% methanol was added to each well, followed by incubation for 30 minutes. After drying, 0.1% crystal violet solution was added. After 30 min, the residual dye was washed away, and 33% acetic acid was added. The OD₅₉₀ was measured to achieve quantitative analysis of biofilm biomass. In the biofilm eradication assay, mature biofilms were formed via cultivation. After the corresponding drugs were added to each group, the biofilms were processed via the crystal violet staining method after 24 h, and three biological replicates were established. Simultaneously, the biofilms were subjected to ultrasonication, and the number of viable bacteria within the biofilms was quantified. Imaging observations were performed via a confocal laser scanning microscope (CLSM; N-STORM, Nikon, Japan). SYTO 9 and PI were used to stain the biofilms, and all imaging parameters (including gain and offset) were kept consistent across all samples.

### 4.6 Membrane integrity assays

In accordance with previous methods[48], the bacterial suspension was adjusted to an OD600 of 0.5. A 10 nmol/L concentration of PI (Beyotime, Shanghai, China) was added to the bacterial suspension, followed by incubation at 37°C for 30 minutes. The suspension was then washed with PBS buffer to remove excess dye. The test drug was added, and the mixture was incubated at 37°C for 2 hours. The fluorescence intensity was measured at Ex535 nm/Em615 nm.

### 4.7 Determination of Reactive Oxygen Species

In accordance with previous methods[49], the bacterial suspension was adjusted to an OD600 of 0.5. A 10 μM DCFH-DA fluorescent probe (Beyotime, Shanghai, China) was added. Then, 100 μL of the stained bacterial suspension and 100 μL of the test drug were added to the wells and incubated for 2 hours, with at least three replicate wells set for each concentration. After incubation, the fluorescence was measured at Ex492 nm/Em525 nm.

### 4.8 Membrane Depolarization Test

In accordance with previous methods[50], the bacterial suspension was adjusted to an OD600 of 0.5. The suspension was pretreated with a 0.5 μM DiSC3(5) probe (Beyotime, Shanghai, China) and incubated at 37°C for 10 minutes. Then, 190 μL of the prestained bacterial suspension was mixed with 10 μL of the test drug and incubated for 2 hours. The fluorescence was measured at Ex622 nm/Em670 nm.

### 4.9 ΔpH measurement

In accordance with previous methods[51], the bacterial suspension was adjusted to an OD600 of 0.5. A 20 μM concentration of BCECF-AM (Beyotime, Shanghai, China) was added. Then, 100 μL of the prestained bacterial suspension was mixed with 100 μL of the test drug and incubated for 2 hours. The fluorescence was measured at Ex500/Em522.

### 4.10 Qualification of ATP

In accordance with previous methods[52], the bacterial suspension was adjusted to an OD600 of 0.5. The test drug was then added, followed by incubation at 37°C for 2 hours. After incubation, the supernatant was collected by centrifugation for subsequent use. The bacterial pellet was resuspended in lysis buffer containing lysozyme and digested to ensure complete cell lysis. The lysate was mixed with an equal volume of luciferase reagent under light-protected conditions. The intracellular ATP content of the bacteria was expressed in relative light units (RLU).

### 4.11 Swimming Motility Assay

*S. suis* SC19 suspension was adjusted to an OD600 of 0.5. A 3 μL aliquot of the bacterial suspension was inoculated at the center of a TSB plate containing 0.3% agar. The plate was incubated at 37°C overnight. The swimming motility of *S. suis* SC19 was observed and photographed, and the maximum migration diameter of the bacterial swimming zone was measured.

### 4.12 Construction of a Mouse Infection Model and therapeutic Efficacy Evaluation

Following activation, strain SC19 was resuspended in sterile PBS to final concentrations of 2.5×10⁸ or 1×10⁸ CFU/mL for two separate experiments. Specific pathogen-free (SPF) female BALB/c mice were intraperitoneally (i.p.) injected with 200 μL of the respective bacterial suspension to establish the infection model. After one hour, the mice received i.p. administration of the drug at specified doses (1, 2, and 4 mg/kg for the survival study; 2 and 4 mg/kg for the bacterial load study). For the survival assay, the mice were monitored for 7 days to record mortality. For the bacterial load assay, the mice were anesthetized at 6 hours post-infection for blood collection via the orbital sinus, and the harvested liver and spleen tissues were homogenized, serially diluted, and plated on TSA medium containing 5% newborn calf serum for enumeration.

All animal experimental schemes and operating techniques were approved by the Animal Experiment Protection, Supervision and Control Committee of Huazhong Agricultural University. The laboratory animal ethics batch number is HZAUMO-2025-0463.

### 4.13 Statistical analysis

Statistical analysis was performed via GraphPad Prism 8.0 software. All the data are presented as the means ± SDs, and comparisons were made via two-tailed unpaired t tests or one-way ANOVA, with statistical significance denoted as *p < 0.05, **p < 0.01, ***p < 0.001, or ns (not significant).

## Authorship contribution statement

Conceptualization: Di Liu and Chang Chen. Supervision: Chen Tan. Methodology: Na Su, Qianqian Bao and Yuhan Zhang. Formal analysis: Xiliang Yang, Wei Fang and Chen Tan. Validation: Na Su and Yuhan Zhang. Writing-original draft: Di Liu. Writing-review & editing: Chenchen Wang and Manli Liu.

## Data availability

Data will be made available on request.

## Institutional Review Board Statement

The animal experiments conformed to the ethical approval of the animal model, and all the experiments were conducted under the guidance of the Protection, Supervision, and Control Committee of Animal Experiments of Huazhong Agricultural University. The study was approved by the Institutional Review Board (or Ethics Committee) of Huazhong Agricultural University (protocol code HZAUMO-2025-0463).

